# Chemo-architecture of area prostriata in adult and developing mice: comparison with presubiculum and parasubiculum

**DOI:** 10.1101/2022.03.01.482588

**Authors:** Sheng-Qiang Chen, Chang-Hui Chen, Xiao-Jun Xiang, Shun-Yu Zhang, Song-Lin Ding

## Abstract

Retrosplenial area 29e, which was a cortical region described mostly in earlier rodent literature, is often included in the dorsal presubiculum (PrSd) or postsubiculum (PoS) in modern literature and commonly used brain atlases. Recent anatomical and molecular studies have revealed that retrosplenial area 29e belongs to the superficial layers of area prostriata, which in primates is found to be important in fast analysis of quickly moving objects in far peripheral visual field. As in primates, the prostriata in rodents adjoins area 29 (granular retrosplenial area), area 30 (agranular retrosplenial area), medial visual cortex, PrSd-PoS, parasubiculum (PaS) and postrhinal cortex (PoR). The present study aims to reveal the chemo-architecture of the prostriata versus PrSd-PoS or PaS by means of a systematic survey of gene expression patterns in adult and developing mouse brains. First, we find many genes that display differential expression across the prostriata, PrSd-PoS and PaS and that show obvious laminar expression patterns. Second, we reveal subsets of genes that selectively express in the dorsal or ventral parts of the prostriata, suggesting the existence of at least two subdivisions. Third, we detect some genes that shows differential expression in the prostriata of postnatal mouse brains from adjoining regions, thus enabling identification of the developing area prostriata. Fourth, gene expression difference of the prostriata from the medial visual cortex and PoR is also observed. Finally, molecular and connectional features of the prostriata in rodents and non-human primates are discussed and compared.

## Introduction

Brodmann appeared to be the first to describe the retrosplenial subarea (area 29e) on the caudal medial aspect of the rabbit brain (Brodmann, 1909). According to him, rabbit area 29e, together with other four retrosplenial subareas (areas 29a-d, or RSg), occupies whole medial surface of the occipital lobe. On his map, rabbit area 29e is a large region labeled among area 29b-d, presubiculum (PrS; or area 27), areas 19 and 20, whereas an agranular retrosplenial area (area 30 or RSag), which was described in other species, is absent in rodents including rabbits (Brodmann, 1909; rabbits belong to lagomorphs in modern literature). Although Brodmann did not illustrate area 29e in brain sections, a conspicuous triangular field located between the dorsal PrS (PrSd, or postsubiculum, PoS) and parasubiculum (PaS) of the rat and mouse brains was treated as the equivalent of area 29e by later scientists (Blackstad, 1956; Haug, 1976; Preston-Ferrer et al., 2016; Slomianka & Geneser, 1991; Vaz Ferreira, 1951) or included in the PrSd-PoS) (e.g., Paxinos & Watson, 2001; Swanson 2018). In many other species, this triangular region was not identified likely because it did not display triangular shape in these species. On the other hand, another unique region, area prostriata (or prostriata), was identified at the location immediately adjoining the striate cortex (i.e., primary visual cortex, V1 or area 17) in human and non-human primates (Allman & Kaas, 1971; Morecraft et al., 2000; Rockland, 2012; Sanides, 1969; Sousa et al., 1991) but not in rodents. In sequential sections stained with Nissl substance, calcium-binding proteins and a non-phosphorylated neurofilament protein, the prostriata in human and non-human primates was found to adjoin the retrosplenial cortex (RS), PrSd-PoS, PaS and posterior cingulate cortex (area 23) at the anterior levels and the V1 and V2 (i.e., secondary visual cortex, or area 18) at the posterior levels (Ding et al., 2003; 2016).

In a survey of the subicular complex across species, the triangular region in rodent brains was also found to adjoin the RS, PrS-PoS, PaS and visual cortex (V1 and V2) (Ding, 2013). This reminds the similar findings from detailed studies of the prostriata in macaque monkeys (Ding et al., 2003; Morecraft et al., 2000). In addition, the triangular region in rodents and the prostriata in non-human primates both receive direct projections from the V1 (Ding, 2013; Sousa et al., 1991). These findings strongly suggest that the triangular region (or area 29e) in rodents is the equivalent of or part of the prostriata in human and non-human primate brains. Our recent studies on this triangular region of the rat and mouse brains have discovered that the triangular region belongs to the superficial layers (layers 2-3) of the prostriata rather than the RS or PrSd-PoS (Chen et al, 2020; Chen et al., 2021; Hu et al., 2020; Lu et al., 2020). These studies have further revealed that rodent prostriata also has deep layers 5 and 6 and a cell-less zone (lamina dissecans), which separates layers 2-3 from 5-6. Although functional studies of area 29e were not carried out previously, our recent findings that the prostriata in rodents receives strong inputs directly from lateral geniculate nucleus (DLG) and V1 (Chen et al., 2021; Lu et al., 2020) support the functions of the prostriata in fast analysis of rapid moving stimuli in far peripheral visual field observed in human and non-human primate brains (Mikellidou et al., 2017; Tamietto and Leopold, 2018; Yu et al., 2012).

In the present study, we aim to systematically investigate the chemo-architecture of the prostriata with focus on laminar and regional differences. This type of information is important for future cell type-specific circuit and functional studies of the prostriata. The second aim is to identify the gene markers for localization of the prostriata at different developmental stages and for developmental study of this region. Our third aim is to compare gene expression patterns of the prostriata and PrSd-PoS or PaS to further illustrate the difference among these structures.

## Materials and Methods

### 1. Animals and in situ hybridization data

In situ hybridization (ISH) raw data were derived from Allen Brain Atlas [ABA, https://mouse.brain-map.org, for adult mouse brains; see Lein et al., 2007] and Allen Developing Mouse Brain Atlas (ADMBA, https://developingmouse.brain-map.org, see Thompson et al., 2014). The ADMBA covers seven ages including four embryonic (postconception days: E11.5, E13.5, E15.5, and E18.5) and three postnatal ages (P4, P14, and P28 days after birth, where day of birth is P0). All experiments were performed in accordance with the Guide for the Care and Use of Laboratory Animals of the Research Ethics Committee. The details for generating these data including probe synthesis, primer design, tissue preparation, condition of hybridization, image processing, and quality control are published online (https://help.brain-map.org/display/mousebrain/Documentation for ABA; https://help.brain-map.org/display/devmouse/Documentation for ADMBA). Briefly, after sectioning, ISH of stains was performed on slides using a semi-automated non-isotopic digoxigenin-labeled colorimetric platform. Riboprobes were labeled with either digoxigenin-UTP or dinitrophenyl-11-UTP (DNP; Perkin Elmer, Waltham, MA, USA). A DNP-labeled probe and a DIG-labeled probe were hybridized simultaneously. Tyramide signal amplification was performed for each probe individually, using either anti-DIG-HRP with tyramide biotin or anti-DNP-HRP with tyramide-DNP for amplification. The sections from embryonic ages were counterstained with nuclear HP Yellow to aid identification of anatomic structures. Finally, it should be mentioned that mRNA expression does not necessarily correlate one-to-one with protein expression but many or at least some of them should correlate. For example, both Calb1 (gene) and calbindin-D28k (the protein coded by Calb1) are strongly expressed in the prostriata, as reported in our previous studies (Chen et al., 2021; Lu et al., 2000).

### 2. Data analysis and image capture

From the ABA and ADMBA portals, we first screened enriched gene expression in the region corresponding to the prostriata defined in our recent studies (Hu et al., 2020; Lu et al., 2020) using the gene finder tool of the Anatomic Gene Expression Atlas (AGEA) application. We then manually examined each selected case to confirm the enriched expression and determine their laminar distribution in the prostriata. Those cases with no obvious expression in the prostriata (except for the cases with clear negative expression in layers 2-3 or 5 of the prostriata, which were labeled as negative marker genes) were excluded from further analysis. The same protocols were used for searching enriched genes in the PrS-PoS and PaS. Finally, the images from the regions of interest were selected and imported into Adobe Photoshop for adjustment of brightness and contrast as well as arrangement and anatomical annotation.

## Results

### 1. Borders, topography and extent of area prostriata

As in our previous studies, we define the boundaries of the prostriata using a combination of cytoarchitecture and molecular markers. To demonstrate the borders, topography and extent of the prostriata in both sagittal and coronal sections, here we use *Pcsk5* as a gene marker for both sectioning planes so that the lamination, shape and extent of the prostriata can be directly visualized and compared on the two planes. Generally, it is relatively difficult to distinguish the prostriata from adjoining regions in Nissl preparations. For example, both the prostriata and PaS display similar cell size and packing density (e.g., Fig. 1a, b). However, some features of the prostriata are still visible such as larger cells in its superficial layers 2-3 compared to adjoining PrSd, RS and V1 (Figs. 1c and the insets in Fig. 1h, i). In *Pcsk5*-ISH stained sagittal sections, the triangular layers 2-3 of the prostriata stand out because of their strong *Pcsk5* expression while no or faint expression is observed in the superficial layers of the neighboring regions such as PaS, PrSd, RS and V1 (Fig. 1d-i). Layers 5 and 6b of the prostriata are also clearly seen since they are *Pcsk5* positive and continue with layers 5 and 6b of adjoining regions, respectively. It is noted the shapes and trajectories of layer 5 of the prostriata vary from lateral (Fig. 1d) to medial (Fig. 1i) sagittal levels. At the most lateral level, layer 5 is straight down into the deep part of the PaS and larger in length than layers 2-3 (Fig. 1d) while at the medial levels layer 5 is slightly curved and smaller in length compared to layers 2-3 in sagittal sections (Fig.1e-i). Compared to surrounding regions, layer 6 of the prostriata is a cell-sparse zone (Fig. 1a-c) and negative for *Pcsk5* (Fig. 1d-f). It is worth mentioning that layers 5 and 6b of the PrSd display stronger and weaker *Pcsk5* expression in comparison with the prostriata.

**Fig. 1.**
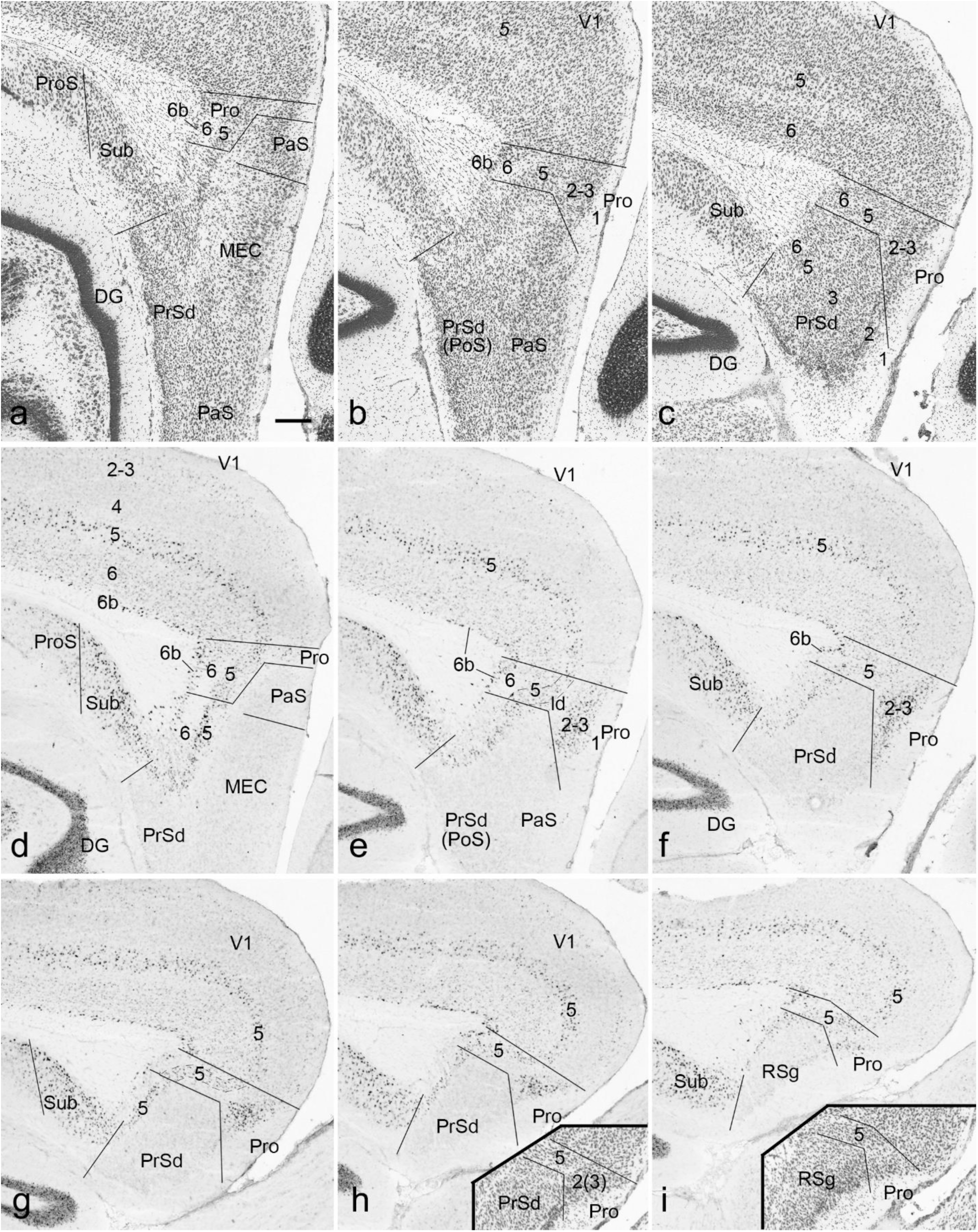
Location, topography and lamination of area prostriata (Pro) in sagittal sections. Arabic numbers indicate cortical layers. (a-c) Nissl-stained sections from three lateral levels (a-c) showing the location and lamination of the Pro and adjoining regions. (d-i) Expression of the gene *Pcsk5* in the Pro from lateral (d) to medial (i) sagittal sections. Layers 2-3 of the Pro is clearly identified with its strong *Pcsk5* expression compared to faint expression in adjoining primary visual cortex (V1) and dorsal presubiculum (PrSd). *Pcsk5* is also expressed in layer 5 of the Pro, V1 and PrSd. The borders of layer 5 in these three regions were determined by layer 5-selective/dominant markers such as *C1ql2* (see Lu et al., 2020), *Lypd1* and *Efnb3* (see Fig. 4). Note the differential shapes and orientation of the layers across different lateral to medial levels. The insets in (h) and (i) show the location of the Pro in Nissl-stained sections adjacent to panels (h) and (i). DG, dentate gyrus; Sub, subiculum; ProS, prosubiculum; PoS, postsubiculum; PaS, parasubiculum; MEC, medial entorhinal cortex; RSg, granular retrosplenial cortex (area 29). Scale bar: 210µm in (a) for all panels.

In *Pcsk5*-ISH stained coronal sections (Fig. 2), both layers 2-3 and 5 of the prostriata can be identified but they look different from those seen in sagittal sections. Most of the RSg does not adjoin the prostriata (Fig. 2a) except its most caudal part (Fig. 2b). From the rostral (Fig. 2b) to caudal (Fig. 2i) levels, the overall sizes of layers 2-3 and 5 change from small (Fig. 2d) to large (Fig. 2e-g) to small (Fig. 2h, i). In coronal sections, the prostriata adjoins the RSg, RSag, RSagl (lateral to RSag), V2M and V1 dorsally, and the PrSd and PaS ventrally (Fig. 2b-i). The most caudal part of the prostriata merges with the postrhinal cortex (PoR) and PaS laterally (Fig. 2i). The PoR strongly expresses *Nnat* while the prostriata, PaS and V1 do not (see the inset in Fig. 2i).

**Fig. 2.**
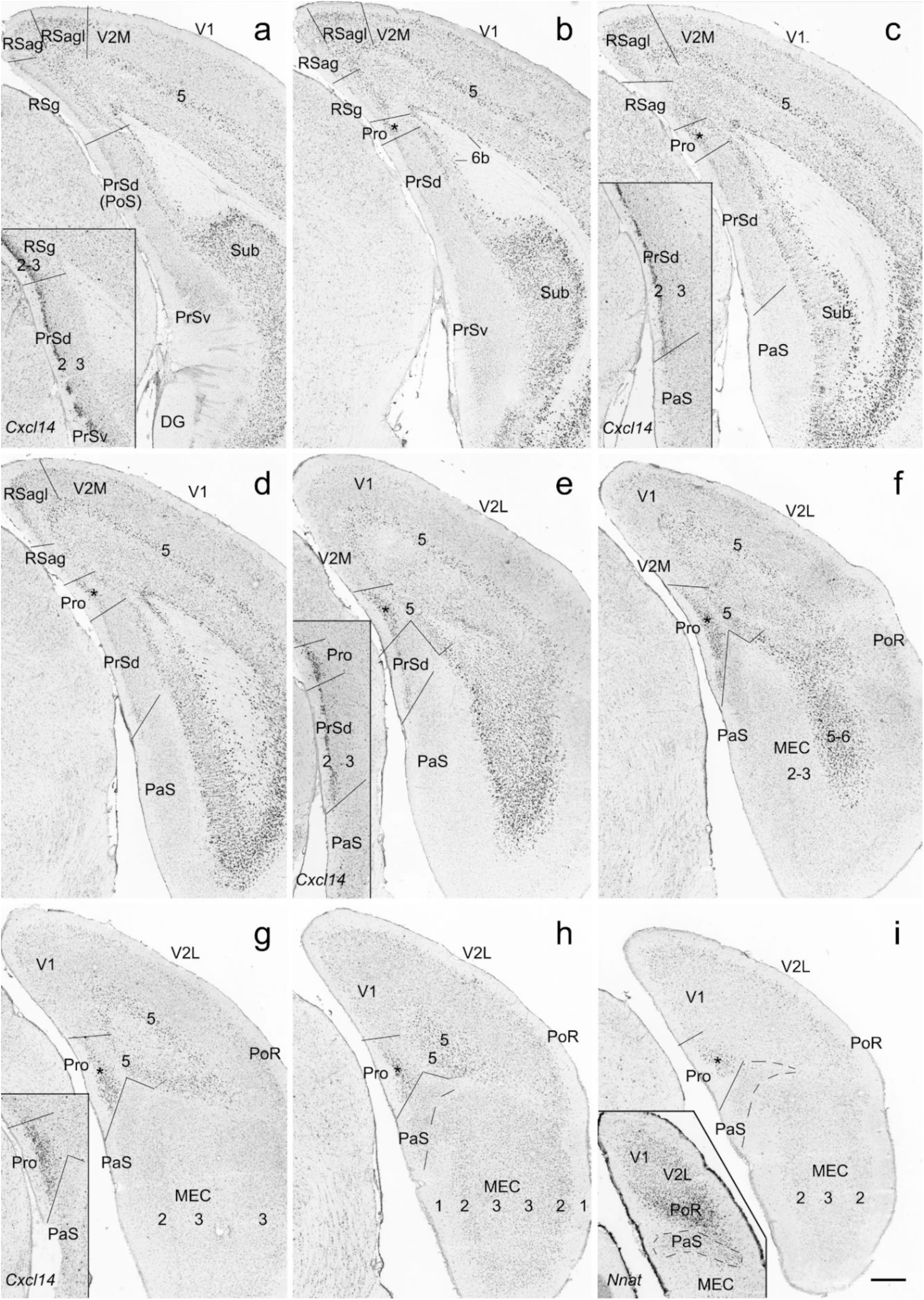
Location, topography and lamination of area prostriata (Pro) in coronal sections. (a-i) *Pcsk5* expression in the Pro from rostral (a) to caudal (i) sections. Layers 2-3 (indicated by star) of the Pro is clearly distinguishable with its strong *Pcsk5* expression compared to no or faint expression in adjoining V1 and PrSd. *Pcsk5* is also expressed in layer 5 of the Pro, RS, V1 and PrSd. The borders of layer 5 with adjoining regions were determined by layer 5-selective/dominant markers such as *C1ql2* (Lu et al., 2020), *Ociad2* and *Slc24a3* (see Fig. 6). Note that the overall sizes, shapes and extent of the Pro vary along rostro-caudal axis with the largest extent at the middle coronal levels (d-g). The insets in (a), (c), (e) and (g) shows the borders between RSg and PrSd (a), PrSd and PaS (c), Pro and PrSd (e), and Pro and PaS (g), respectively, on matched *Cxcl14*-ISH stained sections. *Cxcl14* is strongly expressed in layers 2-3 of RSg, layer 2 of PrSd and layers 2-3 of the Pro while similar layer 2 is not available in the PaS. The inset in (i) is from a *Nnat*-ISH stained section caudal to level (i) displaying the location of the postrihinal cortex (PoR), which strongly expresses *Nnat* in its layers 2-3 (see the inset) and is located caudo-lateral to the Pro and dorsal to the PaS. Layers 2-3 of the Pro and PaS have faint *Nnat* expression. RSag, agranular retrosplenial cortex (area 30); RSagl, lateral agranular retrosplenial cortex; V2M and V2L, medial and lateral secondary visual cortex. Scale bar: 350µm in (a) for all panels.

In *Cxcl14*-ISH stained coronal sections, layers 2-3 of the RSg, layer 2 of the PrSd and layers 2-3 of the prostriata show strong *Cxcl14* expression whereas the PaS is negative (see the insets in Figs. 2a, 2c, 2e and 2g). In fact, layer 2 of the PrSd is composed of densely packed small neurons whereas the PaS does not have this kind of layer 2 and this can be clearly appreciated in Nissl preparations (see Lu et al., 2020).

### 2. Laminar gene markers for layers 2-3, 5 and 6 of area prostriata

Based on the definition and location of the prostriata, the expression of many genes is found to be enriched or restricted in layers 2-3 with no or faint expression in other layers of the prostriata. These genes include *Igsf3, Inpp4b, Prkca, Rxfp1, Ntsr1, Homer2, Tcerg1l, Fstl1, Cxcl14, Nr2f1, Cdc42ep3, Otof, Unc5d, Syt10, Gabrb3, Car4, Spns2, Palmd, Fat3* and *Cux2* (Fig. 3a-t) as well as *Slc17a6, Col23a1, Adra2a, Adra2c, Htr1b, Htr1f* and *Ntsr1* (not shown). Gene expression enriched or restricted in layer 5 is also observed and these genes include *Parm1, Nnat, Fezf2, Lypd1, Efnb3, Col6a1, Pex5l, Nos1ap* (Fig. 4a-h), *Crym* (Fig. 4j), *Gpr161* and *Ntng1* (not shown). Gene markers for layer 6b include *Tle4, Crym, Sdk2, Limch1* (Fig. 4i-l), *Drd1, Drd5, Chrna4, Chrna5* and *Adra2a* (not shown). Interestingly, the expression of many marker genes for layer 6 of the neocortex does not extend into the prostriata and these genes include *Col6a1, Nos1ap, Tle4, Crym, Sdk2* and *Limch1* (Figs. 4f, i-l). It should be noted that the PrSd-PoS has less expression of *Parm1, Nnat, Fezf2, Lypd1, Efnb3, Col6a1, Pex5l, Nos1ap* (Fig. 4a-h) and *Crym* (Fig. 4j) in layer 5 compared to the prostriata.

**Fig. 3.**
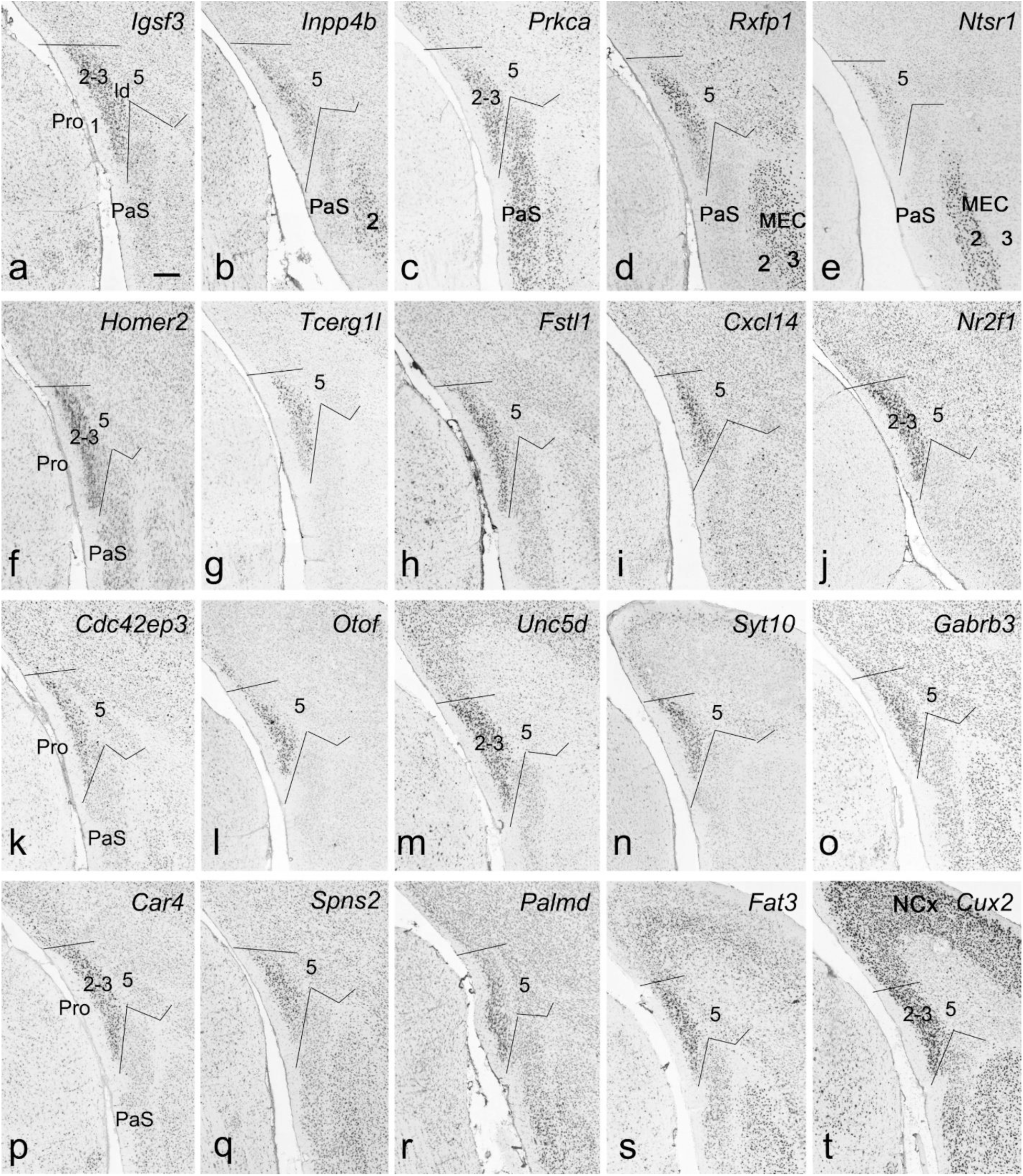
Gene expression in layers 2-3 of the prostriata (Pro). (a-t) Examples of 20 genes enriched in layers 2-3 of the Pro in coronal sections. Gene names are indicated on top-right corners of each panel. It is clear that *Igsf3, Inpp4b, Prkca, Rxfp1, Ntsr1, Homer2, Tcerg1l, Fstl1, Cxcl14, Nr2f1, Cdc42ep3, Otof, Unc5d, Syt10, Gabrb3, Car4, Spns2, Palmd and Fat3* (a-s) are enriched in layers 2-3 of the Pro but not the PaS (except *Prkca* in c). Strong expression of *Inpp4b* in layer 2 of medial entorhinal cortex (MEC; b), *Prkca* in layers 2-3 of the PaS (c), *Rxfp1* in layer 3 of the MEC (d), *Ntsr1* in layer 2 of the MEC (e), *Homer2* in layers 2-3 of the PaS (f) and *Cux2* in layers 2-4 of the neocortex (NCx; t) are also observed. Scale bar: 210µm in (a) for all panels.

**Fig. 4.**
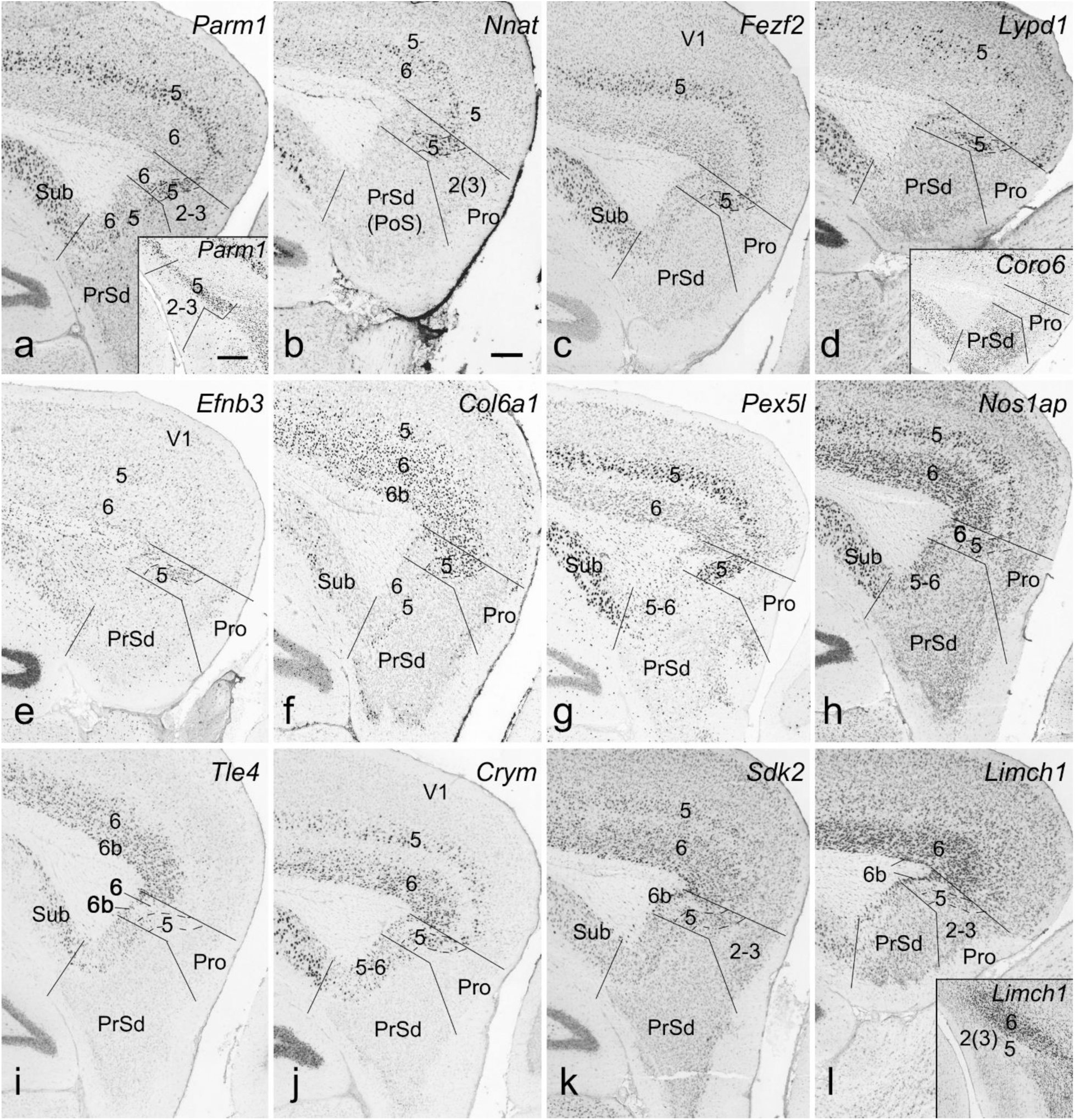
Gene expression in layers 5-6 of the prostriata (Pro). Gene names are indicated on top-right corners of each panel; layer 5 is outlined by dashed lines. (a-g) Examples of 7 genes enriched in layer 5 of the Pro in sagittal sections. It is clear that *Parm1, Nnat, Fezf2, Lypd1, Efnb3, Col6a1 and Pex5l* (a-g) are enriched in layer 5 of the Pro but not the PrSd. Note the strong expression of these genes in layer 5 of the overlying V1 with exception of *Efnb3*, which has much fewer expression in V1 (e). Inset in (a) shows *Parm1* expression in layer 5 of the Pro in a coronal section. Inset in (d) shows faint and strong *Coro6* expression in the Pro and PrSd, respectively, in a matched sagittal section. (h-l) Examples of 5 genes enriched in layer 6b of the Pro in sagittal sections. Strong expression of *Nos1ap* (h) and *Crym* (j) is also seen in layer 5 of the Pro. Inset in (l) shows *Limch1* expression in layer 6 of the overlying V1 in a coronal section. Note that layer 6 of the Pro is a cell-sparse zone with less gene expression. Scale bars: 210µm in (b) for all panels; 350µm in the inset in (a) for all insets.

### 3. Negative gene markers for layers 2-3 and 5-6 of area prostriata

Some genes are not expressed in layers 2-3 or 5-6 of the prostriata but display strong expression in neighboring regions. This pattern is also helpful in distinguishing the prostriata from adjoining PrSd and V1. *Prkcb* (Fig. 5a-d) and *Pcp4* (Fig. 5e-h) are the negative marker genes for layers 2-3 and 5 of the prostriata, respectively, in sharp contrast to layers 2-3 and 5 of the PrSd and V1. Another example is the expression of *Coro6* (see the inset in Fig. 4d), *Ier5* and *Cdh8* (not shown) in layer 5 but not in layers 2-3 and 6 of the prostriata; these genes are strongly expressed in layers 2-3 of adjoining PrSd or V1. Final example for the difference is the expression of *Loc434300* (Fig. 5i-l), *Trib2, Nts, Nr4a2* and *Col25a1* (not shown), which are negative and positive in layer 5 of the prostriata and PrSd, respectively.

**Fig. 5.**
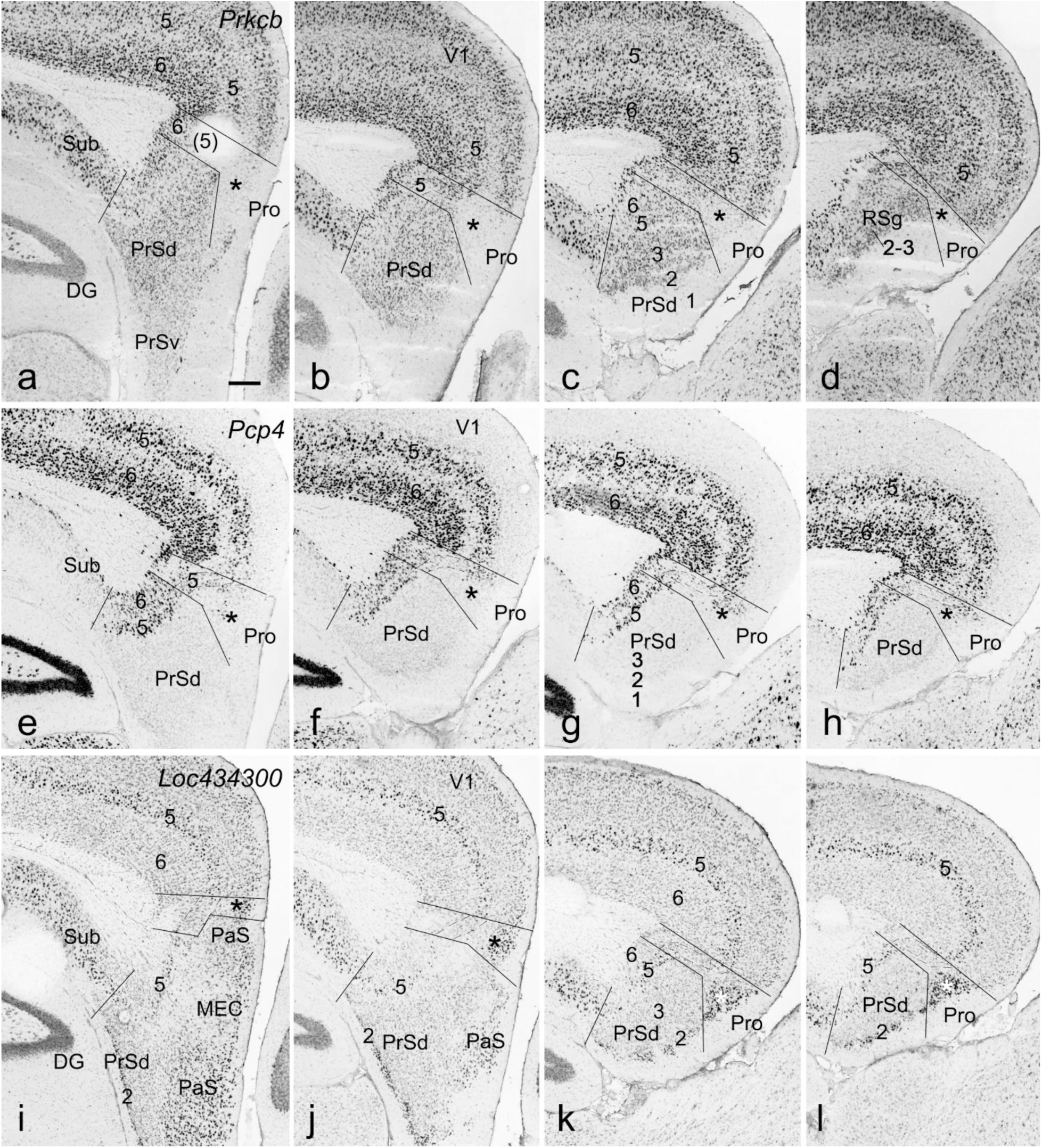
Negative gene expression in layers 2-3 and 5 of the prostriata (Pro). All panels are from sagittal sections. Layers 2-3 of the Pro are indicated by the stars. (a-d) Negative *Prkcb* expression in layers 2-3 of the Pro from lateral (a) to medial (d) levels. Note the surrounding regions with strong *Prkcb* expression. (e-h) Negative *Pcp4* expression in layer 5 of the Pro from lateral (e) to medial (h) levels. In contrast to the Pro, layer 5 of the adjoining PrSd and V1 strongly express *Pcp4*. Note the weaker *Pcp4* expression in the dorsal part of layers 2-3 of the Pro. (i-l) *Loc434300* expression in the Pro from lateral (i) to medial (l) levels. Negative and strong *Loc434300* expression is found in layers 5 and 2-3 of the Pro, respectively. Note that layer 5 in both PrSd and V1 expresses this gene. Strong expression is also visible in the PaS and layer 2 of the PrSd. Scale bar: 210µm in (a) for all panels.

### 4. Gene expression enriched in both layers 2-3 and 5 of area prostriata

In addition to the layer-specific gene expression, other genes are expressed in both layers 2-3 and 5. Figure 6 shows 16 genes with this expression pattern. Among these genes, *Gria3, Lifr, Ssbp2, Syt17, Icam5, Id2, Smpd4, Ociad2, Tmem150c* (Fig. 6a-i) and *Spon1* display higher expression in layers 2-3 than in layer 5 while some other genes such as *Chst8, Gda, Slc24a3, Kcnn2, 6430573f11Rik, Mal2* and *Cntn4* (Fig. 6j-p), *Bcl6, Lamp5, Drd2, Adrbk2, Chrm3* (not shown) show similar expression in both layers 2-3 and 5. Finally, still other genes such as *Gpr123* have stronger expression in layer 5 than in layers 2-3 (not shown).

**Fig. 6.**
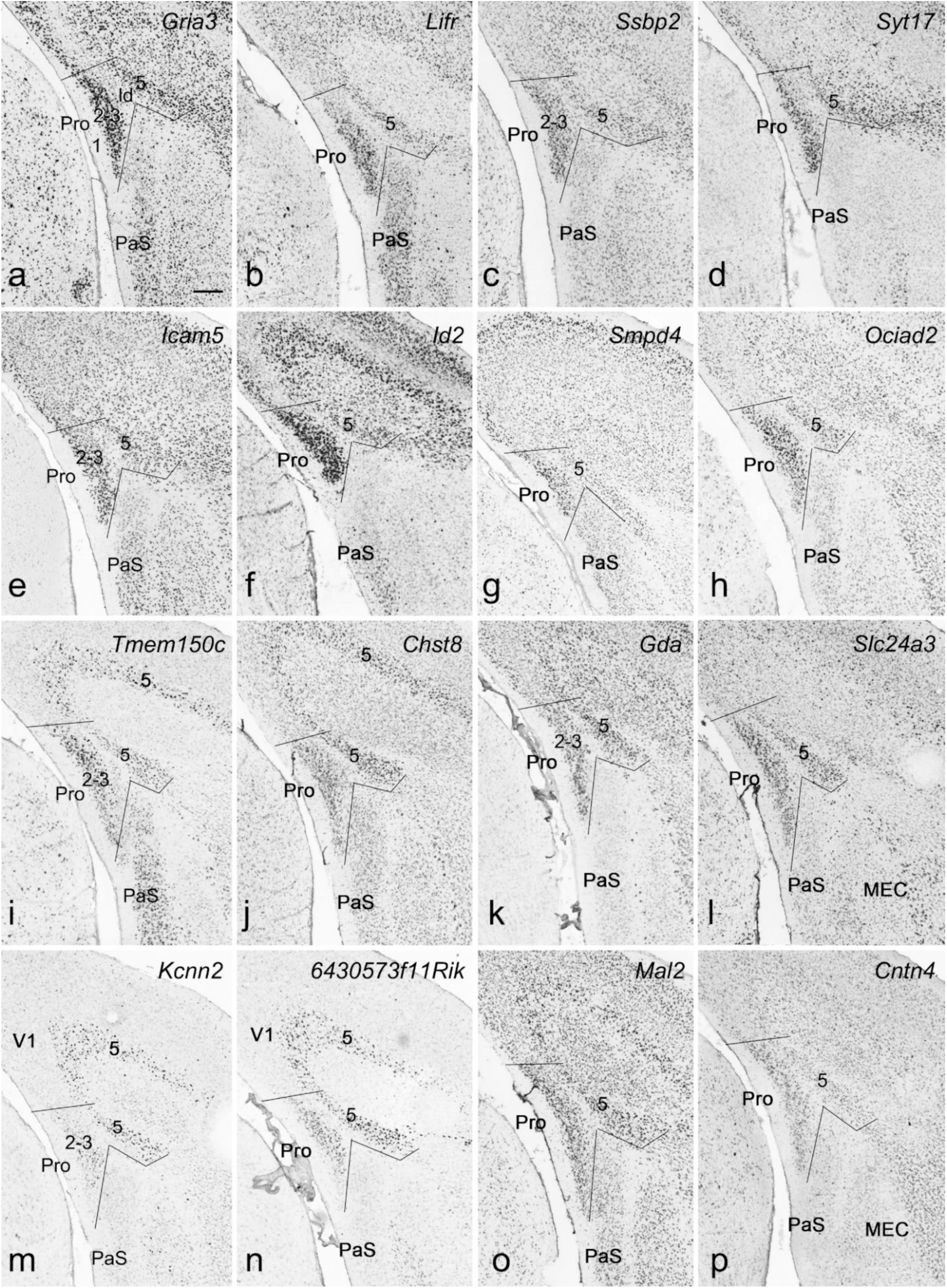
Gene expression in both layers 2-3 and 5 of the prostriata (Pro). All panels are from coronal sections. Gene names are indicated on top-right corners of each panel. (a-i) Examples of 9 genes with strong expression in layers 2-3 and relatively weaker expression in layer 5 of the Pro. These genes are *Gria3, Lifr, Ssbp2, Syt17, Icam5, Id2, Smpd4, Ociad2 and Tmem150c* (a-i). Note also the strong expression of *Gria3* (a), *Lifr* (b), *Icam5* (e) and *Tmem150c* (i) in the PaS. (j-p) Examples of 7 genes with similar expression intensity in both layers 2-3 and 5 of the Pro. These genes are *Chst8, Gda, Slc24a3, Kcnn2, 6430573f11Rik, Mal2* and *Cntn4* (j-p). Scale bar: 210µm in (a) for all panels.

### 5. Subdivisions of area prostriata

Since the prostriata in macaque monkeys was divided into anterior and posterior subareas (Pro-a and Pro-p) based on different cyto- and chemo-architecture with Pro-p adjoining the V1 (Ding et al., 2003), we have searched the ABA to see if evidence for differential gene expression exists within mouse prostriata. As shown in the sequential sagittal sections, *Col12a1* is clearly expressed in the dorsoposteior part (Pro-d) but not in the ventroanterior part (Pro-v) of the prostriata although the dorsoventral difference is difficult to be appreciated in Nissl-stained sections (Fig. 7a-j). In contrast, *Penk* expression in the prostriata displays an opposite pattern (Fig. 7k-o). Sometimes, regional gene expression is not seen throughout all levels of the prostriata. For instance, *Scnn1a* is mostly expressed in the Pro-d at the lateral levels (Fig. 7p, q) with no or less at the medial levels (Fig. 7r, s). At more medial levels where the RSg appears, Scnn1a is also expressed in layer 5a of the RSg (Fig. 7t). In our survey, many genes are found to express in the Pro-d and these genes include *Col12a1, Col5a1, Chn2, Rorb, Scnn1a, Loc433288, 2900026a02rik, Spag5, Whrn, Flrt2, Sytl2, Tmem150c, Plcxd2, Rims3* (Fig. 8a-n) and *Pcp4* (Fig. 5e-h) as well as *Kitl, Zmat4, Cacna1g, Adra1d, Chrna7, Trib2* and *Igfbp5* (not shown). In contrast, much fewer genes are observed to express only in the Pro-v and these genes include *Penk* (Fig. 7p, q), *Cdh24, Gpr88* (Fig. 8o, p) and *Pcdh8* (not shown).

**Fig. 7.**
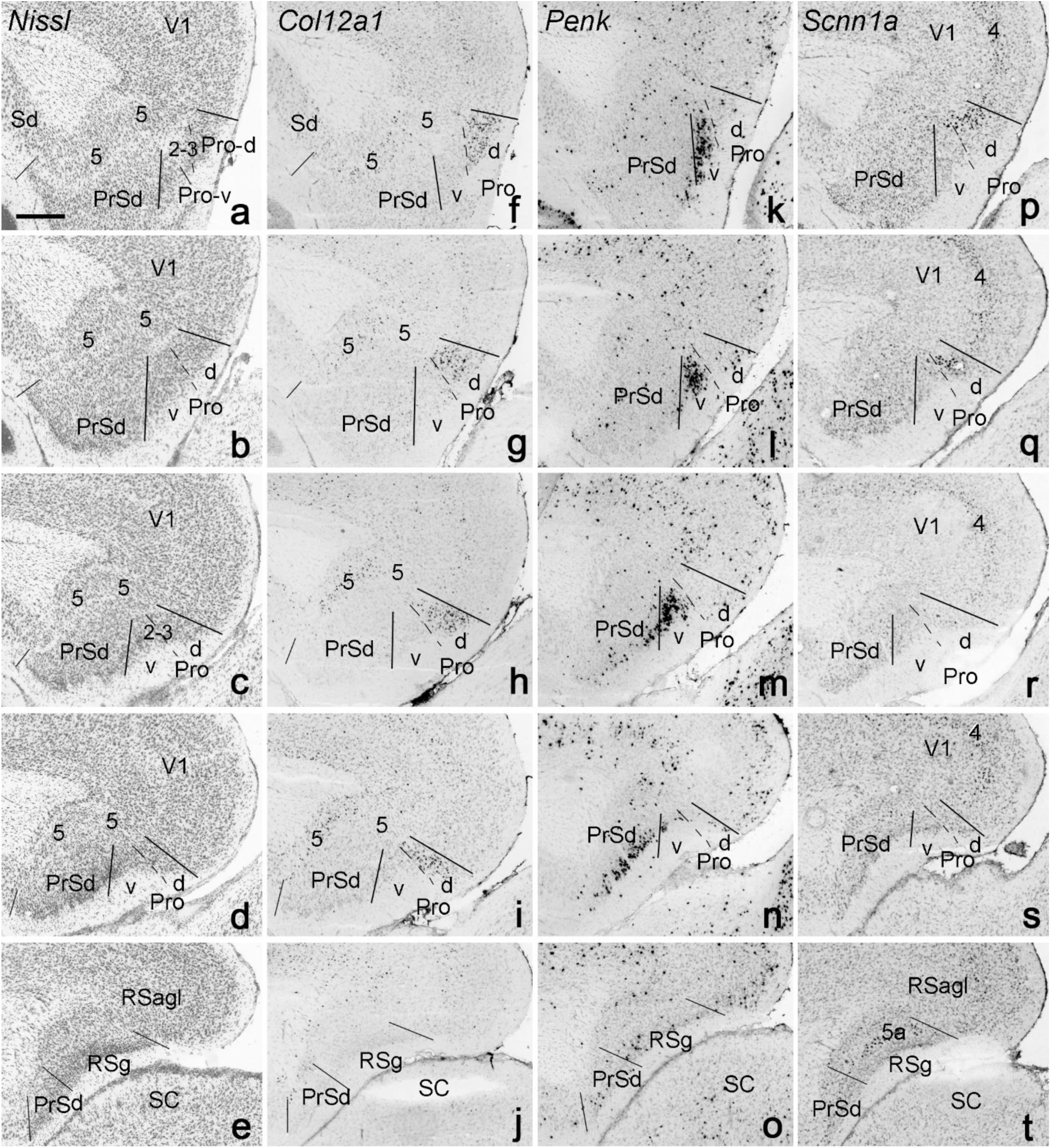
Identification of two subdivisions of the prostriata (Pro) in sequential sagittal sections. Lateral (top) to medial (bottom) sections are arranged along each column. (a-e). Locations of the Pro in sequential Nissl-stained sections. The two subdivisions (Pro-d and Pro-v) of the Pro are not well distinguishable in these sections and a granular layer 4 does not appear to exist in both subdivisions. (f-j). *Col12a1* expression in layers 2-3 of the Pro-d in sequential sections adjacent to (a-e), respectively. (k-o). *Penk* expression in layers 2-3 of the Pro-v in sequential sections. Note also the strong *Penk* expression in layer 2 of the medial PrSd (n). (p-t). *Scnn1a* expression in deep layer 3 of the Pro-d in sequential sections. Note that *Scnn1a* expression in the Pro is only seen at the lateral levels (p, q) rather than the medial levels (r, s). Scale bars: 300µm in (a) for (a-t).

**Fig. 8.**
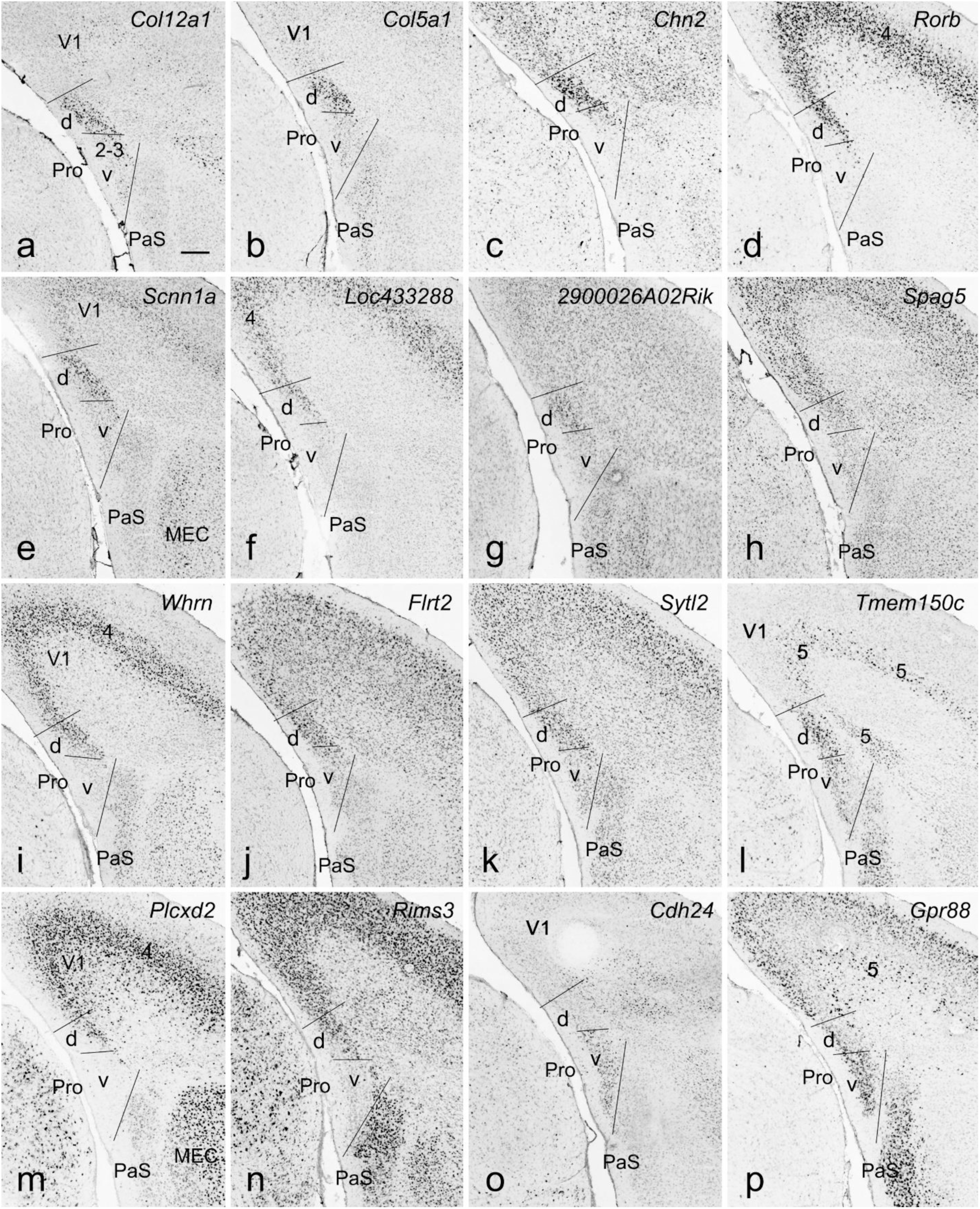
Differential gene expression in two subdivisions of the prostriata (Pro). All panels are from coronal sections. Gene names are indicated on top-right corners of each panel. (a-n) Examples of 14 genes enriched in layers 2-3 of the dorsal subdivision (Pro-d). Note that some genes enriched in layer 4 of the V1 (e.g., *Rorb, Whrn* and *Plcxd2*) also express in the Pro-d (d, i, m). (o-p) Examples of two genes enriched in layers 2-3 of the ventral subdivision (Pro-v). An additional example of weaker *Pcp4* expression in Pro-d of the Pro is shown in sagittal sections in Figure 5e-h. Scale bar: 210µm in (a) for all panels.

### 6. Localization of area prostriata in developing mouse brains

To identify the prostriata in developing mouse brains, we have explored the ADMBA and found that the prostriata is not recognizable in the prenatal brains at E11.5, E13.5, E15.5 and E18.5 (not shown). However, some genes are clearly expressed in the postnatal prostriata from P4 to P28. As early as P4, for example, weaker expression of *6430573f11rik* is observed in layers 2-3 and 5 of the prostriata while strong expression of this gene is seen in layer 5 of the PrSd (Fig. 9a-d). Genes *Htr1a, Id2, Fstl1* (not shown) and *Nr2f1*(Fig. 9e-h) display a different expression pattern with strong and weaker expression in layers 2-3 and 5-6, respectively. *Nr2f1* shows moderate expression in layer 5 of the PrSd and strong expression in the visual cortex (V1) (Fig. 9e-h). Finally, weaker *Tcerg1l* expression can be found in layers 2-3 and 5 of the prostriata compared to strong expression in layer 5 of the V1 and layer 2 of the PrSd (Fig. 9e-h). Stronger expression of *Etv1, Grp* and *Crym* in layer 5 of the prostriata versus PrSd also makes the prostriata stand out (not shown). It is worth mentioning that layer 5 of the prostriata at P4 does not express *C1ql2*, which is strongly expressed in adult prostriata (see Lu et al., 2020). At P14, strong *6430573f11rik* expression is detected in layers 2-3 and 5 of the prostriata while in the PrSd its expression is contained in layer 5 with no expression in layers 2-3 (Fig. 10a-d) and this pattern remains from P14 onward to adult (not shown). Strong *C1ql2* expression is detected and restricted in layer 5 of the prostriata with no expression in adjoining V1 and PrSd at P14 (Fig. 10e-g) and P28 (e.g., Fig. 10h). Additionally, *Tcerg1l* expression is mostly found in layers 2-3 of the prostriata, layer 2 of the PrSd and layer 5 of the V1 from P14 (Fig. 10i-l) to adult (not shown). Finally, *Nr2f1* is strongly expressed in layers 2-3 of the prostriata with no or faint expression in the PrSd from P14 onward (not shown). The negative expression of *Neurod6* and *Cdh8* in layers 2-3 of the prostriata also makes the prostriata identifiable from the neighboring regions such as PrSd, V1 and PaS at P4 (Fig. 11a-d for sagittal; 11e-h for coronal sections), P14 (Fig. 11i-l) and P28 (not shown). Taken together, the prostriata can be reliably identified in postnatal mouse brains using gene markers such as those described above.

**Fig. 9.**
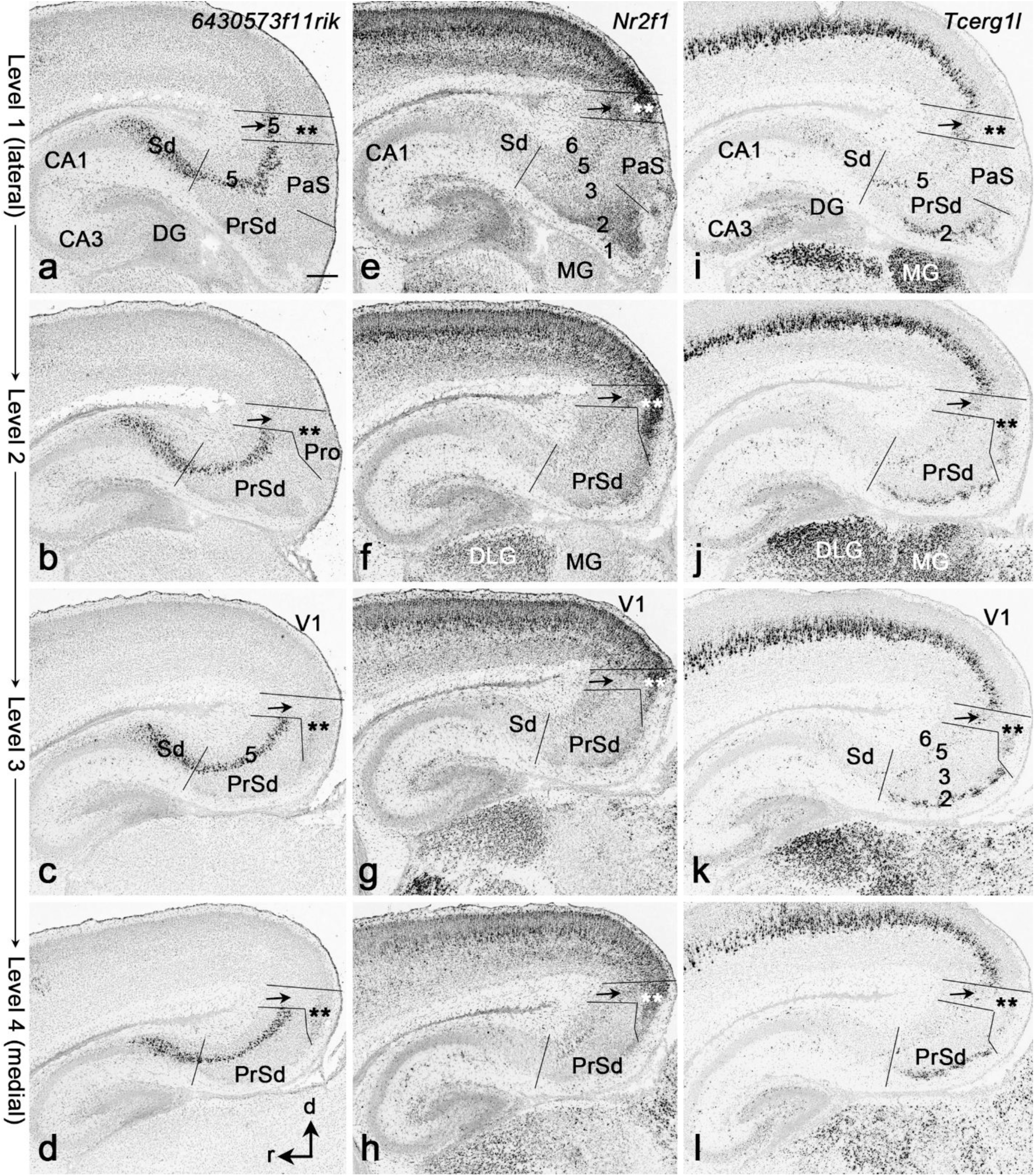
Identification of the prostriata (Pro) in neonate mice (P4). All panels are from sagittal sections. Lateral (top) to medial (bottom) sections at levels 1-4 are arranged along each column. (a-d) Weak *6430573f11rik* expression in both layers 5 (arrows) and 2-3 (stars) of the Pro. In contrast, strong expression is detected in dorsal subiculum (Sd) and layer 5 of the PrSd. Note that the expression of this gene is not yet detected in layer 5 of the V1 at P4. (e-h) Strong *Nr2f1* expression in layers 2-3 of the Pro (stars) with weaker expression in layer 5 (arrows). Note that this gene is expressed in dorsal lateral geniculate nucleus (DLG) but not in medial geniculate nucleus (MG). (i-l) Weak *Tcerg1l* expression in layers 2-3 and 5 of the Pro. Note the strong expression in layer 2 of PrSd and layer 5 of the V1. It should be mentioned that no expression of *C1ql2* was observed in layer 5 of the Pro at P4 and thus not shown here. Scale bar: 220µm in (a) for (a-l).

**Fig 10.**
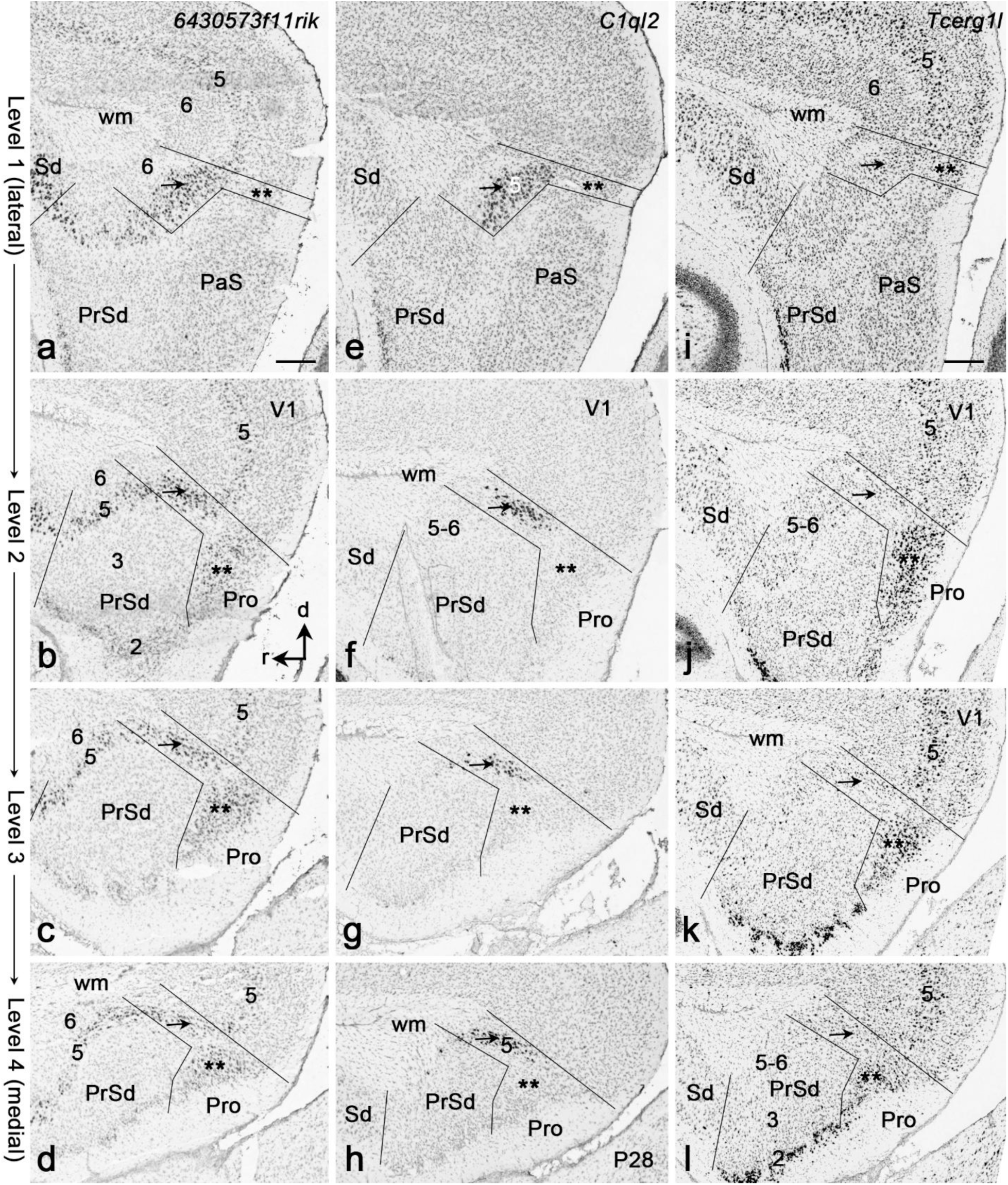
Identification of the prostriata (Pro) at P14 and P28. All panels are from sagittal sections. Lateral (top) to medial (bottom) sections at levels 1-4 are arranged along each column. (a-d) *6430573f11rik* expression in both layers 5 (arrows) and 2-3 (stars) of the Pro at P14. This gene is also expressed in dorsal subiculum (Sd) and layer 5 of the V1 and PrSd. (e-h) *C1ql2* expression in layer 5 of the Pro (arrows) at P14 (e-g) and P28 (h). Note that layer 5 in neighboring regions is negative for *C1ql2*. (i-l) *Tcerg1l* expression in layers 2-3 of the Pro (stars) at P14. Note the strong expression in layer 2 of the PrSd (k, l). Strong and weaker expression is seen in layer 5 of the V1 and Pro, respectively. Scale bars: 210µm in (a) for (a-h); 220µm in (i) for (i-l).

**Fig 11.**
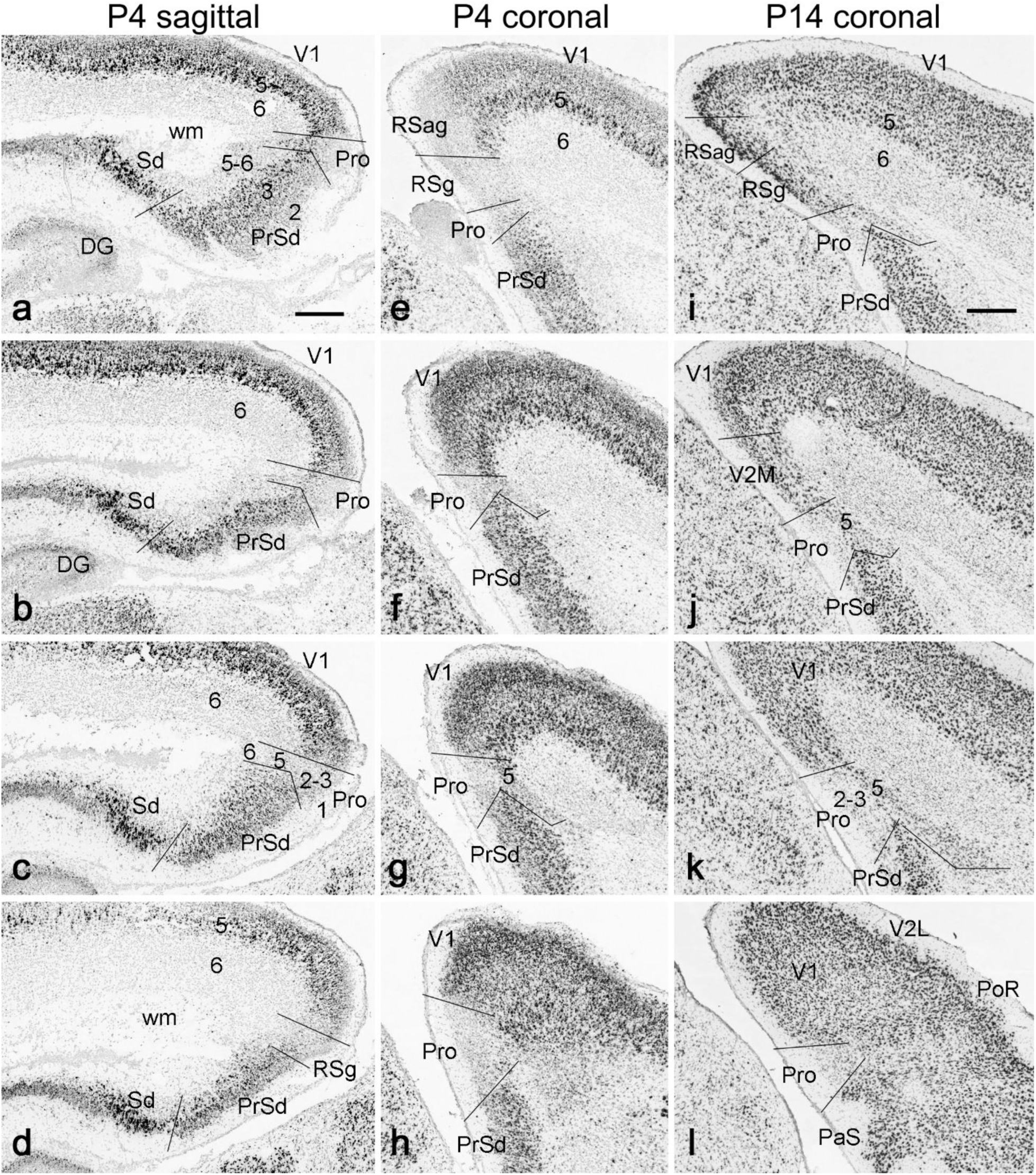
Identification of the prostriata (Pro) using the negative marker gene *Cdh8*. (a-h) Negative expression of *Cdh8* in layers 2-3 of the Pro at P4 in sequential sagittal (a-d, from lateral to medial levels) and coronal (e-h, from rostral to caudal levels) sections. (i-l) Negative expression of *Cdh8* in layers 2-3 of the Pro at P14 in sequential coronal sections (i-l is from rostral to caudal levels). Note the weaker *Cdh8* expression in layer 5 of the Pro. In contrast to the Pro, strong *Cdh8* expression is obvious in adjoining V1 and PrSd. Scale bars: 250µm in (a) for (a-h); 300µm in (i) for (i-l).

### 7. Comparison of area prostriata with presubiculum and parasubiculum

The genes with differential expression in the prostriata versus PrS-PoS includes *Pcsk5, 6430573f11rik, Dgkb, Scn4b* and *Prkca* in layers 2-3 (Figs. 1, 2, 12a-b, 12f-g), and *Nnat, Fezf2, lypd1, Efnb3, Col6a1, Pex5l, Pcp4, Loc434300, C1ql2, Lxn and Deptor* in layers 5-6 (Figs. 4a-I, 5e-l,10e-h, 12c-e). The genes with distinct expression in the prostriata versus PaS include *Pcsk5, Igsf3, Inpp4b, Rxfp1, Ntsr1, Tcerg1l, Fstl1, Cxcl14, Nr2f1, Cdc42ep3, Otof, Unc5d, Syt10, Syt17, Ociad2, Gda, Kcnn2, 6430573f11rik, Cntn4, Prkca* and *Fn1* (Figs. 2, 3, 6, 12), in addition to *Wfs1* (Luuk et al., 2008; Ding, 2013; Chen et al., 2020). Differential expression of *Pcsk5, Prkca* and *Fn1* is also clearly seen in the PrS versus PaS (Figs. 2c-e,12g,12l).

**Fig 12.**
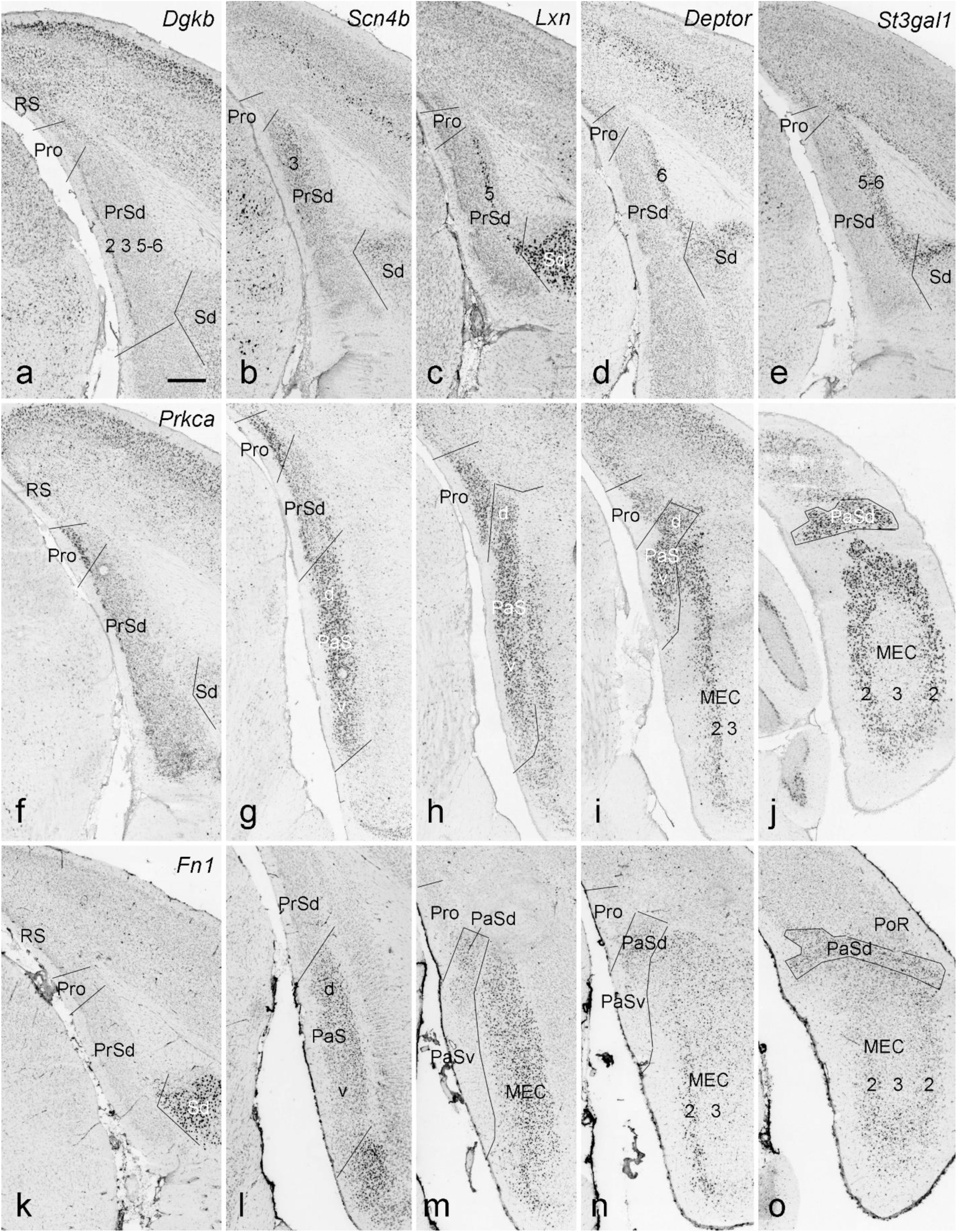
Molecular markers for the presubiculum and parasubiculum. (a-e) Examples of molecular markers for layers 1 (a), 2 (b), 5 (c), 6 (d) and 5-6 (e) of the dorsal presubiculum (PrSd). Gene names are indicated on the top of each penal. Note that the expression of these genes does not extend into the prostriata (Pro). (f-j) Differential expression of *Prkca* in the Pro, PrSd and PaS from rostral (f) to caudal (j) coronal levels. The PrSd shows relatively weaker expression than adjoining Pro and PaS, which displays strong expression. Strong *Prkca* expression is also seen in layer 2 of the medial entorhinal cortex (MEC). (k-o) Differential expression of *Fn1* in the PrSd and PaS from rostral (k) to caudal (o) coronal levels. No *Fn1* expression is detected in the Pro and PrSd, whereas strong and faint *Fn1* expression is observed in the dorsal (PaSd) and ventral PaS (PaSv), respectively. Scale bar: 350µm in (a) for all panels.

At postnatal ages (P4, P14 and P28), some genes such as *Ddit4l, Nnat* and *Wfs1* display differential expression in the prostriata versus PrS-PoS or PaS. Specifically, *Ddit4l* is strongly expressed in the superficial layers of the PrS-PoS and PaS but only faintly in the prostriata. *Nnat* strongly expresses in the superficial layers of PaS rather than the prostriata and PrS-PoS, in which the expression is found in the deep layers. Finally, while no expression exists in the PrS-PoS, *Wfs1* is expressed strongly and weakly in the PaS and prostriata, respectively.

### 8. Comparison of area prostriata in the marmosets and mice

To explore if the prostriata has some conservative gene expression between mice and marmoset monkeys, we have carried out a survey on the Riken marmoset database (https://gene-atlas.brainminds.riken.jp), which mainly contains the ISH data from postnatal stages (mostly at P0; see Kita et al., 2021; Shimogori et al., 2018). As expected, some genes indeed show similar expression patterns between the two species. For example, in the neonate marmosets (P0), weak and strong *Id2* expression is observed in layers 2-3 of area 29 (A29 in Fig. 13a, b) and the Pro (Fig. 13b, c), respectively. In addition, the superficial layers of areas 30, 23 and V1 also display strong *Id2* expression (Fig. 13a-d). V1 in marmosets has a very thick layer 4 (negative for *Id2*; see Fig. 13c, d), making it distinguishable from the prostriata, where almost no granular layer 4 is detected. Another gene *Grp* is mainly expressed in layers 2-3 of A29 (Fig. 13e, f) and layer 5 of many cortical regions including the Pro (Fig. 13e-h). The *Grp* expression in layer 5 is stronger in the Pro compared to adjoining regions (Fig. 13f, g). In the neonate mice (P4), strong *Id2* expression exists in layers 2-3 of both A29 (not shown) and the Pro (Fig. 13i), and this pattern remains at P14 (not shown). However, weak and strong *Id2* expression (similar to the marmosets) is detected in area 29 and the Pro at P28. The *Grp* expression pattern in the neonate mice (P4) is generally similar to that in the marmosets. Specifically, *Grp* is strongly expressed in layers 2-3 of area 29 (Fig. 13j) and layer 5 of the Pro (k).

**Fig 13.**
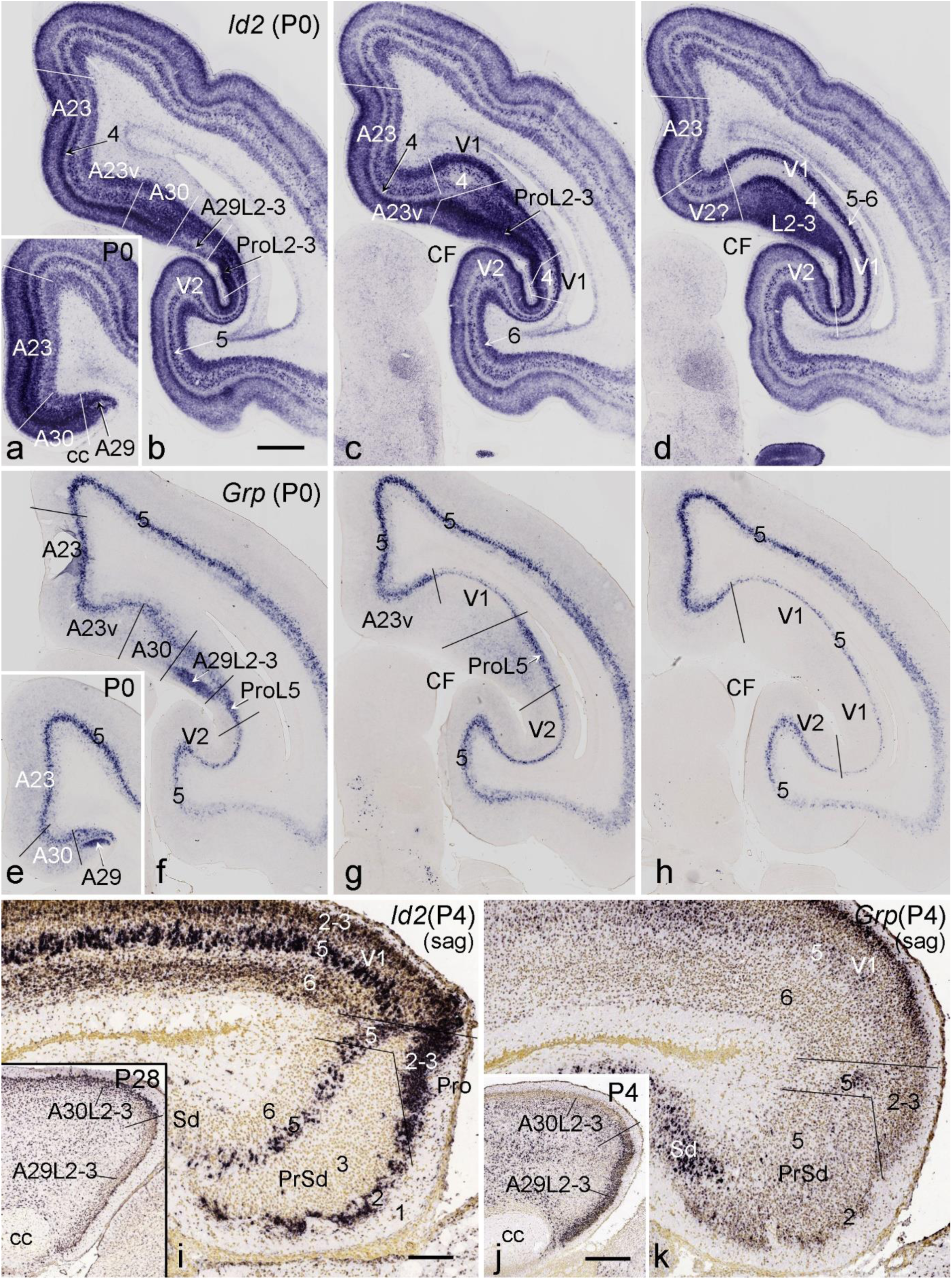
Comparison of *Id2* and *Grp* expression in the prostriata (Pro) of the marmosets and mice. Panels (a-d) and (e-h) are four sequential coronal sections from the marmoset at P0. (a-d) *Id2* expression in the Pro, area 29 (A29), area 30 (A30), area 23 (A23), V1 and V2 of the marmoset. Note the weak and strong *Id2* expression in layers 2-3 of A29 (a, b) and the Pro (b, c), respectively. The superficial layers of A30, A23 and V1 also display strong *Id2* expression. (e-h). *Grp* expression in the Pro, A29, A30, A23, V1 and V2 of the marmoset. Strong *Grp* expression in layers 2-3 of A29 (e, f) and relatively stronger expression in layer 5 of the Pro (f, g) compared to adjoining regions are observed. Layer 5 of the V1 shows the weakest *Grp* expression (g, h). (i-k) *Id2* (i) and *Grp* (j, k) expression in the Pro, PrSd, V1, A29 and A30 in sagittal sections from the mice at P4. Strong *Id2* expression is observed in layers 2-3 of the Pro and layers 2 and 5 of the PrSd (i), while strong *Grp* expression is seen in layers 2-3 of A29 (j) and layer 5 of the Pro (k). The inset in (i) shows *Id2* expression in A29 and A30 of a mouse at P28. At this age, weak and strong *Id2* expression exists in A29 and A30, respectively, and this pattern is similar to that found in the marmoset (a, b). It should be mentioned that strong *Id2* expression exists in both A29 and A30 at P4 and P14 (data not shown). corpus callosum, cc; CF, calcarine fissure. Scale bars: 1mm in (a) for (a-h); 200µm in (i) for (i) and (k); 420µm in (j) for (j) and the inset in (i).

## Discussion

The present study has revealed clear differential expression of many genes in the prostriata and adjoining structures such as PrS, PaS, RS, PoR and V1. The distinct gene expression strongly supports the prostriata as an independent anatomical entity (Ding, 2013; Lu et al., 2020) rather than as part of RSg (area 29e) (Blackstad, 1956; Haug, 1976; Slomianka and Geneser, 1991; Vaz Ferreira, 1951) or part of PrS-PoS (Paxinos and Watson, 2001; Swanson 2018). The similar topographic relationship of the prostriata with neighboring regions (see Figs. 1, 2) compared to non-human primates (Morecraft et al., 2000; Ding et al., 2003; Fig. 13) further supports its nature as the equivalent of the primate prostriata (Allman and Kaas, 1971; Ding et al., 2003; Morecraft et al., 2000; Paxinos et al., 2012; Rockland, 2012; Sanides, 1969; Sousa et al., 1991). Similar expression patterns of some conservative genes such as *Id2* and *Grp* in the prostriata of the neonate mice and marmosets (Fig. 13) also confirm the rodent homolog of the prostriata. In this study, we have discovered many layer-specific gene expression or cell types in the prostriata and subdivided the prostriata into two main parts based on differential molecular signature. We have also delineated the location and extent of the prostriata in postnatal mouse brains. All these findings would provide a fundamental base for future cell-type related structural, functional and developmental exploration of this unique region.

### 1. Laminar organization and cell types of area prostriata

Like in human and non-human primate brains, the prostriata in rodent brains is a limbic cortex characterized by lack of a clear granular layer 4, where a lamina dissecans appears to separate the superficial layers 2-3 and deep layers 5-6 (Ding, 2013; Hu et al., 2020; Lu et al., 2020). In the present study, we have revealed layer-specific expression of many genes in the prostriata. These include specific expression in layers 2-3 or layer 5 or both, indicating the existence of different cell types in the prostriata. These findings are consistent with our recent connectional studies in the rat and mouse brains (Chen et al., 2020; Chen et al., 2021; Hu et al., 2020). For example, layers 2-3 project mainly to contralateral layers 2-3 (Chen et al., 2020) and ipsilateral PrS-PoS (Chen et al., 2021) while layer 5 projects mainly to the lateral dorsal thalamic nucleus (LD), ventral lateral geniculate nucleus (VLG), pulvinar (Pul), zona incerta (ZI), pretectal nucleus (PTN) and pontine nucleus (PN) (Chen et al., 2021). Many afferent projections to the prostriata also display obvious laminar distribution, indicating different inputs innervating distinct layers or cell types in the prostriata (Chen et al., 2021; Hu et al, 2020). Interesting future studies would be functional correlation of these laminar and/or cell type specific connections via generation of cell type specific Cre-mouse lines using the gene markers revealed in the present study.

### 2. Subdivisions of area prostriata

Based on gene expression difference within the prostriata we have divided the prostriata into the dorsal and ventral parts (Pro-d and Pro-v; see Figs. 7, 8). Pro-d adjoins RS and visual cortex (V2M and V1) dorsally while Pro-v abuts PrSd-PoS and PaS ventrally. Pro-d displays no or incipient granular layer 4 and thus corresponds to Pro-p (i.e., posterior part) of the prostriata in macaque monkeys (Ding et al., 2003). Pro-v shows a clear lamina dissecans and thus corresponds to Pro-a (i.e., anterior part) of the macaque prostriata. In this study, many genes are observed to express in layers 2-3 of the Pro-d but not Pro-v (Fig. 8). Interestingly, afferent projections from the rostral midline thalamic region and rostral DLG heavily innervate layers 2-3 of the Pro-d but not Pro-v in rodent (Chen et al., 2021; Hu et al., 2020). Cortical projections from the V1 also tend to target the Pro-d more heavily than the Pro-v (Lu et al., 2020). In contrast to Pro-d, we find only a limited number of genes that express in layers 2-3 of the Pro-v although many genes are enriched in both Pro-d and Pro-v. Detailed circuit and functional correlation of these two prostriata subareas remains to be investigated.

### 3. Development of area prostriata

No studies were previously carried out on the development of the prostriata in any species. To localize the prostriata in developing mouse brains we have searched the large dataset, ADMBA (see Thompson et al., 2014) and find that the mouse prostriata is not identifiable prenatally. However, starting from P4 onward, the prostriata can be reliably identified using some positive gene markers such as *6430573f11rik, Nr2f1, Tcerg1l* and *Id2*, as well as some negative markers such as *Cdh8, Prkcb* and *Pcp4*. Additionally, our recent pilot study on the connectivity of the prostriata in postanal rats has revealed the projections from the prostriata to medial V1 at P7 and 14 (unpublished data). It worth noting that, in a recently published prenatal human brain atlas, the prostriata is also not recognizable at PCW 15 and 21 (Ding et al., 2022) although it is possible to identify it at later stages due to the long period of prenatal development. Our localization of the prostriata in the postnatal mouse brains would provide an anatomical base for future studies related to prostriata development and functional maturation.

### 4. Functional consideration of area prostriata

Although the prostriata region in rodent brains has been treated as a part of area 29e or PrS-PoS since Brodmann (Brodmann, 1909), its function has not been investigated and has long been considered as a mystery. On the other hand, the prostriata in non-human primates, first described in later 60s (see Sanides, 1969), was reported to be responsive to visual stimuli (Cuenod et al., 1965; Rosa et al., 1997) but no detailed studies have been carried out until recently. A relatively recent study of the marmoset monkeys has revealed that prostriata neurons have enormous receptive fields and short response latencies and are selective to stimulus speed rather than direction of motion (Yu et al., 2012). A more recent study has found, using population receptive field mapping, that the prostriata in human brains responds strongly to very fast motion in the far peripheral field (Mikellidou et al., 2017; Tamietto and Leopold, 2018). These findings suggest that the prostriata is very important in monitoring far peripheral visual field for unexpected fast-moving objects, particularly dangerous ones. However, the underlying neural circuits and mechanisms for these functions remain to be explored (Rockland, 2012). The discovery of rodent equivalent of the prostriata enables direct investigation of brain-wide circuits of the prostriata (Ding, 2013; Lu et al., 2020). Our recent efforts in mouse and rat brains have revealed that the prostriata receives direct projections from rostral DLG and medial V1 and projects directly to the effectors responsible for visuomotor behaviors such as LD, PTN, PN, VLG, Pul and ZI (Chen et al., 2021; Lu et al, 2020). Such short visual pathways avoid multiple synaptic delay and would provide the neural substrates for fast processing of rapid moving stimuli in the far peripheral visual field. Additionally, strong commissural connections between two sides of the prostriata (Chen et al., 2020) would enable quick integration of related information from two sides of peripheral visual fields. The rodent prostriata also projects to other regions such as visual and auditory association cortex (V2 and A2), limbic cortex (orbitofrontal, retrosplenial, postrhinal, ectorhinal and entorhinal cortices) and subcortical regions such as caudal putamen and claustrum although these projections are usually weak (Chen et al., 2021; unpublished data). Consistent with the rodents, the prostriata projections to the primary visual (Sousa et al, 1991), secondary auditory (Falchier et al., 2010) and orbitofrontal (Barbas, 1993) cortices were reported in non-human primates although the data about other projections mentioned above are not available. On the other hand, the primate prostriata was reported to project to middle temporal area (Palmer & Rosa, 2006; Rosa et al., 1993), frontal pole (Burman et al., 2011) and rostral cingulate motor cortex (Morecraft et al., 2000), whereas these regions are not yet defined in rodent brains.

### 5. Comparison of the prostriata with presubiculum, parasubiculum and area 23

Although the rodent prostriata has strong reciprocal connections with the PrS-PoS (Chen et al., 2020; 2021), many major connectional differences exist between the two structures in addition to the molecular differences described in the present study. First, the PrS-PoS receives direct projections from hippocampal CA1 (Ding et al., 2020) while the prostriata does not (Chen et al., 2021; Hu et al., 2020). Second, the prostriata but not the PrS-PoS receives strong projections directly from V1 and DLG. Third, the PrS-PoS has strong projections to lateral mammillary nucleus (mouse: Ding, 2013; rat: Yoder & Taube, 2011) whereas the prostriata does not project to this nucleus (Chen et al., 2021). Fourth, the PrS-PoS rather than the prostriata projects strongly to both sides of the medial entorhinal cortex (MEC; Chen et al., 2020 for mouse; van Groen & Wyss, 1990 & Honda et al., 2008 for rat). Finally, the prostriata projects strongly to PTN, PN, VLG, Pul and ZI (Chen et al., 2021) while the PrS-PoS does not.

In addition to distinct gene expression patterns shown in the present study, the PaS also displays differential connections from the prostriata. For instance, the PaS but not the prostriata receives strong inputs from basolateral nucleus of amygdala (mouse: Ding, 2013; rat: van Groen & Wyss, 1990) and parafascicular-centromedian thalamic nucleus (unpublished data). On the other hand, the PaS does not receive projections from the visual and auditory cortices and DLG while the prostriata does (Chen et al., 2021). As for the efferent projections, the PaS projects strongly to the superficial layers of the MEC bilaterally (Chen et al., 2020 & Hu et al. 2020 for mouse; Caballero-Bleda & Witter, 1993 and van Groen & Wyss, 1990 for rat) whereas the prostriata displays weak projections to the deep layers of the MEC (Chen et al., 2021). Additionally, as mentioned above, the PaS does not project to PTN, PN, VLG, Pul and ZI, which are the major targets of the prostriata (Chen et al., 2021).

Previous studies in non-human primates showed that area 23 (A23) also adjoins the prostriata (Ding et al., 2003; Paxinos et al., 2012). This indicates that A23 (including A23v; see Fig. 13) does not belong to the two subdivisions of the prostriata (Ding et al., 2003) although one study (Palmer and Rosa 2006) suggested that A23v in the marmosets could be equated to one of the two subdivisions proposed by Ding et al. (2003). Whether rodents have a homologous A23 remains a mystery because A23 has not been reported so far in rodents. In our recent studies, we have found that the three subdivisions of the rodent RS (RSg, RSag and RSagl) display differential gene expression (Hu et al. 2020; Lu et al, 2020). Since RSg and RSag respectively correspond to areas 29 and 30, it is possible that RSagl or part of it corresponds to A23 in terms of spatial topography of these three subdivisions (see Fig. 13a). However, A23 in non-human primates has a thin but clear layer 4 (Ding et al., 2003; Fig. 13b, c), whereas RSagl in rodents has almost no layer 4 (Hu et al. 2020; Lu et al, 2020). Similarly, the posterior subdivision (Pro-p) of the prostriata in the macaque monkeys has a very thin layer 4, while both subdivisions of the rodent prostriata do not appear to have a layer 4 (see Fig. 7). Therefore, multimodal data including connectional and comparative ones are needed to establish the correspondence of A23 between these two species and the relationship between the prostriata and A23.

Functionally, the head direction cells, grid cells, and border cells were reported to exist in the PrS-PoS (Boccara et al., 2010; Preston-Ferrer et al. 2016; Taube et al. 1990) as well as in the PaS (Tang et al., 2016). These results indicate that the PrS-PoS and PaS are mainly involved in spatial processing and navigation (Dalton & Maguire, 2017; Preston-Ferrer et al. 2016; Sanguinetti-Scheck & Brecht, 2020). However, it is currently not known if those cells or other specialized cells exist in the region corresponding the prostriata. Taken together, the prostriata in rodents shows many different anatomical, molecular and functional features from the PrS-PoS and PaS and thus should be treated as a distinct entity as in human and non-human primates.

## Abbreviations and full names of the genes in this study

2900026a02rik: RIKEN cDNA 2900026A02 gene
6430573f11rik: RIKEN cDNA 6430573F11 gene
Adra1d: adrenergic receptor, alpha 1d
Adra2a: adrenergic receptor, alpha 2a
Adra2c: adrenergic receptor, alpha 2c
Adrbk2: adrenergic receptor kinase, beta 2
Bcl6: B cell leukemia/lymphoma 6
C1ql2: complement component 1, q subcomponent-like 2
Cacna1g: calcium channel, voltage-dependent, T type, alpha 1G subunit
Car4: carbonic anhydrase 4
Cdc42ep3: CDC42 effector protein (Rho GTPase binding) 3
Cdh8: cadherin 8
Cdh24: cadherin 24
Chn2: chimerin 2
Chrm3: cholinergic receptor, muscarinic 3, cardiac
Chrna4: cholinergic receptor, nicotinic, alpha polypeptide 4
Chrna5: cholinergic receptor, nicotinic, alpha polypeptide 5
Chrna7: cholinergic receptor, nicotinic, alpha polypeptide 7
Chst8: carbohydrate (N-acetylgalactosamine 4-0) sulfotransferase 8
Cntn4: contactin 4
Col5a1: collagen, type V, alpha 1
Col6a1: collagen, type VI, alpha 1
Col12a1: collagen, type XII, alpha 1
Col23a1: collagen, type XXIII, alpha 1
Col25a1: collagen, type XXV, alpha 1
Coro6: coronin 6
Crym: crystallin, mu
Cux2: cut-like homeobox 2
Cxcl14: chemokine (C-X-C motif) ligand 14
Ddit4l: DNA-damage-inducible transcript 4-like
Deptor: DEP domain containing MTOR-interacting protein
Dgkb: diacylglycerol kinase, beta
Drd1: dopamine receptor D1
Drd2: dopamine receptor D2
Drd5: dopamine receptor D5
Efnb3: ephrin B3
Etv1: ets variant 1
Fat3: FAT tumor suppressor homolog 3 (Drosophila)
Fezf2: Fez family zinc finger 2
Flrt2: fibronectin leucine rich transmembrane protein 2
Fn1: fibronectin 1
Fstl1: follistatin-like 1
Gabrb3: gamma-aminobutyric acid (GABA) A receptor, subunit beta 3
Gda: guanine deaminase
Gpr88: G-protein coupled receptor 88
Gpr123: G protein-coupled receptor 123
Gpr161: G protein-coupled receptor 161
Gria3: glutamate receptor, ionotropic, AMPA3 (alpha 3)
Grp: gastrin releasing peptide
Homer2: homer homolog 2 (Drosophila)
Htr1a: 5-hydroxytryptamine (serotonin) receptor 1A
Htr1b: 5-hydroxytryptamine (serotonin) receptor 1B
Htr1f: 5-hydroxytryptamine (serotonin) receptor 1F
Icam5: intercellular adhesion molecule 5, telencephalin
Id2: inhibitor of DNA binding 2
Ier5: immediate early response 5
Igfbp5: insulin-like growth factor binding protein 5
Igsf3: immunoglobulin superfamily, member 3
Inpp4b: inositol polyphosphate-4-phosphatase, type II
Kcnn2: potassium intermediate/small conductance calcium-activated channel, subfamily N, member 2
Kitl: kit ligand
Lamp5: lysosomal-associated membrane protein family, member 5
Lifr: leukemia inhibitory factor receptor
Limch1: LIM and calponin homology domains 1
Loc433288: gene Loc433288
Loc434300: gene Loc434300
Lxn: latexin
Lypd1: Ly6/Plaur domain containing 1
Mal2: mal, T cell differentiation protein 2
Neurod6: neurogenic differentiation 6
Nnat: neuronatin
Nos1ap: nitric oxide synthase 1 (neuronal) adaptor protein
Nr2f1: nuclear receptor subfamily 2, group F, member 1
Nr4a2: nuclear receptor subfamily 4, group A, member 2
Ntng1: netrin G1
Nts: neurotensin
Ntsr1: neurotensin receptor 1
Ociad2: OCIA domain containing 2
Otof: otoferlin
Palmd: palmdelphin
Parm1: prostate androgen-regulated mucin-like protein 1
Pcdh8: protocadherin 8
Pcp4: Purkinje cell protein 4
Pcsk5: proprotein convertase subtilisin/kexin type 5
Penk: preproenkephalin
Pex5l: peroxisomal biogenesis factor 5-like
Plcxd2: phosphatidylinositol-specific phospholipase C, X domain containing 2
Prkca: protein kinase C, alpha
Prkcb: protein kinase C, beta
Rims3: regulating synaptic membrane exocytosis 3
Rorb: RAR-related orphan receptor beta
Rxfp1: relaxin/insulin-like family peptide receptor 1
Scn4b: sodium channel, type IV, beta
Scnn1a: sodium channel, nonvoltage-gated 1 alpha
Sdk2: sidekick homolog 2 (chicken)
Slc17a6: solute carrier family 17 (sodium-dependent inorganic phosphate cotransporter), member 6
Slc24a3: solute carrier family 24 (sodium/potassium/calcium exchanger), member 3
Smpd4: sphingomyelin phosphodiesterase 4
Spag5: sperm associated antigen 5
Spns2: spinster homolog 2
Spon1: spondin 1, (f-spondin) extracellular matrix protein
Ssbp2: single-stranded DNA binding protein 2
Syt10: synaptotagmin X
Syt17: synaptotagmin XVII
Sytl2: synaptotagmin-like 2
Tcerg1l: transcription elongation regulator 1-like
Tle4: transducin-like enhancer of split 4, homolog of Drosophila E(spl)
Tmem150c: transmembrane protein 150C
Trib2: tribbles homolog 2 (Drosophila)
Unc5d: unc-5 homolog D (C. elegans)
Wfs1: Wolfram syndrome 1 homolog (human)
Whrn: whirlin
Zmat4: zinc finger, matrin type 4

## Data Availability Statement

All datasets presented in this study are included in the article. These data are available online (http://www.brain-map.org) and from the corresponding author upon reasonable request.

## Ethics Statement

The animal study was reviewed and approved by the Institutional Animal Care and Use Committee of the Allen Institute.

## Author Contributions

Conceptualization: SLD; data investigation and analysis: SLD, SQC, CHC, XJX, SYZ; supervision and writing: SLD. All authors have read and approved the submitted manuscript.

## Funding

This work is partially supported by National Natural Science Foundation of China (#31771327).

## Conflict of Interest

The authors declare that they have no conflicts of interest.

## Acknowledgments

The authors would like to thank Allen Institute for providing publicly accessible mouse brain *in situ* hybridization dataset.

## Notes

### Competing Interest Statement

The authors have declared no competing interest.

